# Limitations of renal arterial hemodynamic measures as candidate biomarkers of renal sympathetic innervation

**DOI:** 10.64898/2026.09.15.751794

**Authors:** Carson Dettmer, Alicia M. Schiller, Irving H. Zucker, Yiannis S. Chatzizisis, Han-Jun Wang, Peter Ricci Pellegrino

## Abstract

**Introduction:** The key role of maladaptive renal sympathetic activation in hypertension has led to the development of catheter-based renal denervation therapies. Unfortunately, the lack of practical physiological biomarkers impairs patient selection and prevents intraprocedural feedback for renal denervation.

**Methods:** We tested whether renal arterial hemodynamic measures and the beat-to-beat variability of these measures reflect renal sympathetic innervation in two tightly controlled preclinical models of renal denervation. Bilateral renal hemodynamic data were obtained from ten conscious unilaterally denervated rabbits and ten anesthetized unilaterally denervated pigs that subsequently underwent stepwise catheter-based radiofrequency denervation of the contralateral kidney. Renal arterial mechanics were assessed by quantification of wave speed and input impedance modulus and phase shift. Variability was quantified as the within-recording standard deviation of each metric when measured on a beat-to-beat basis.

**Results:** Wave speed trended down after surgical denervation in rabbits (P = 0.054) but not swine (P = 0.57); wave speed variability was unchanged in both models. Renal arterial input impedance modulus was not significantly affected by surgical denervation. Surgical renal denervation increased input impedance phase shift in rabbits (P = 0.0048) but not in swine (P = 0.85). The beat-to-beat variability of modulus did not differ significantly between innervated and surgically denervated kidneys in either species. Surgical renal denervation decreased phase shift variability in swine (P = 0.0048) but not rabbits. The difference in phase shift variability between kidneys in swine was eliminated after a single round of branch-vessel ablation (P = 0.42).

**Discussion:** Renal arterial wave speed, input impedance, and their beat-to-beat variability did not consistently or dose-dependently reflect renal sympathetic innervation in these models. These findings argue against their use as direct physiological surrogates for renal sympathetic outflow.

## Introduction

The sympathetic nervous system is a master regulator of homeostasis and a critical pathophysiological mediator of diseases like hypertension, heart failure, and chronic kidney disease (Scott-Solomon, Boehm and Kuruvilla, 2021). In preclinical hypertensive models and hypertensive patients, the renal sympathetic nerves elevate blood pressure through sympatho-excitatory afferent signaling and efferent effects on renin release, sodium handling, and renal vascular control (DiBona and Kopp, 1997; Foss, Fink and Osborn, 2016). Renal denervation has thus emerged as a therapeutic intervention for patients with treatment-resistant hypertension (Kiuchi *et al*., 2019).

A major barrier to renal denervation is the lack of biomarkers to guide patient selection beforehand and confirm effective denervation during the procedure (Esler, 2014; Paton *et al*., 2026). Renal denervation has demonstrated antihypertensive efficacy in contemporary randomized trials, but treatment response remains heterogeneous and the procedure still lacks a practical physiological endpoint of technical success (Azizi *et al*., 2018, 2022; Böhm *et al*., 2020; Mahfoud *et al*., 2022; Saxena *et al*., 2022). A clinically practical marker of renal sympathetic activity could improve both selection of patients most likely to benefit and confidence that adequate denervation has been achieved.

Renal sympathetic nerve activity dynamically regulates renal arterial vascular tone (Schiller, Pellegrino and Zucker, 2016). However, these vascular effects are not well-captured by conventional hemodynamic measurements because renal autoregulatory mechanisms maintain mean renal blood flow in the face of physiological perturbations (Carlström, Wilcox and Arendshorst, 2015).

Moreover, sympathetic outflow is inherently time-varying, raising the possibility that temporal fluctuations in vascular function may contain physiological information lost by time-averaged hemodynamic measures (Guild *et al*., 2001).

Arterial hemodynamic properties provide a complementary way to interrogate vascular physiology because they characterize the dynamic pressure-flow relationship rather than mean flow alone. The Framingham Heart Study showed that pulse wave velocity is a predictive biomarker for cardiovascular event risk (Mitchell *et al*., 2010). In the coronary circulation, measures like the instantaneous wave-free ratio capture physiology that predict the functional significance of coronary stenoses (Davies *et al*., 2017).

Arterial wave speed quantifies the speed at which pressure waves propagate along an artery and is influenced by arterial stiffness and vascular tone (Davies *et al*., 2006). Preliminary clinical reports have suggested that renal arterial wave speed may predict the blood pressure response to catheter-based renal denervation (Finegold *et al*., 2016). Whether wave speed directly reflects renal sympathetic innervation, however, remains uncertain.

Input impedance describes the frequency-dependent relationship between pulsatile arterial pressure and blood flow in terms of both modulus, an index of magnitude, and phase shift, an index of timing (O’Rourke, 1982). Augmentation of sympathetic outflow to the pulmonary circulation by electrical stimulation of the stellate ganglion produces increases in pulmonary artery input impedance modulus at stimulation frequencies that do not affect pulmonary arterial resistance (Pace, 1971). Whether sympathetic innervation affects renal artery input impedance has not been established.

To address these questions, we leveraged two tightly controlled animal models of renal sympathetic denervation: conscious rabbits with chronic unilateral surgical denervation and swine with unilateral surgical denervation followed by stepwise contralateral catheter-based denervation. We sought to identify arterial hemodynamic properties that consistently reflected surgical renal denervation across both experimental models and, in swine, demonstrated a dose-response relationship with progressive catheter-based denervation. Because sympathetic outflow is inherently dynamic, we evaluated both absolute values and beat-to-beat variability of wave speed and input impedance. We hypothesized that renal denervation would decrease renal artery wave speed and input impedance modulus and decrease variability of these measures over time.

## Materials and Methods

### Experimental Overview

This study was a secondary analysis of previously acquired hemodynamic datasets from two published models of renal sympathetic denervation: a rabbit model of chronic unilateral surgical denervation and a swine model of unilateral surgical denervation followed by stepwise contralateral catheter-based radiofrequency denervation (Pellegrino *et al*., 2020, 2025) . The present analysis focused on renal arterial hemodynamic properties derived from those datasets, including wave speed, input impedance, and their temporal variability. All procedures were reviewed and approved by the University of Nebraska Medical Center Institutional Animal Care and Use Committee and carried out in accordance with the NIH Guide for the Care and Use of Laboratory Animals.

### Rabbit Model

Detailed surgical procedures and denervation validation have been reported previously (Pellegrino *et al*., 2020). In brief, ten adult male New Zealand White rabbits (3.3-4.2 kg) underwent unilateral surgical renal denervation, with the contralateral kidney serving as an innervated internal control.

Under general anesthesia, rabbits were instrumented with an arterial pressure telemeter in the abdominal aorta and transit-time volumetric blood flow probes on both renal arteries. After a two-week postoperative recovery period, arterial pressure and bilateral renal blood flow were recorded in conscious animals. Completeness of unilateral denervation was functionally evaluated using the nasopharyngeal reflex as described previously.

### Swine Model

Detailed descriptions of the surgical and percutaneous procedures and denervation validation have been reported previously (Pellegrino *et al*., 2025). In brief, ten adult male swine (44-59 kg) underwent unilateral surgical renal denervation, with the contralateral kidney serving as an innervated control. After a seven-day recovery period, each pig underwent repeat study under ketamine-midazolam anesthesia, during which percutaneous femoral arterial access was obtained for simultaneous measurement of bilateral renal arterial blood flow velocity and abdominal aortic pressure. The innervated kidney subsequently underwent stepwise catheter-based radiofrequency renal denervation, with hemodynamic measurements repeated after each round of ablation. Ablations proceeded from the largest renal branch artery to the remaining branch arteries, followed by the distal and proximal main renal artery.

### Wave Speed Analysis

Renal arterial wave speed was calculated using the single-point sum-of-squares method from simultaneous arterial pressure and renal flow signals (Davies *et al*., 2006). Wave speed was very sensitive to artifacts in the arterial pressure waveform, and thus all beats with peak |dP/dt| > 1.5 times the median peak |dP/dt| across all beats in the recording were discarded, resulting in the exclusion of 76 out of 76,123 cardiac cycles across all recordings (0.10%). Wave speed was then calculated using all remaining beats across the entire recording to obtain a time-averaged estimate. To characterize temporal behavior, wave speed was also calculated separately for each included cardiac cycle. In swine, renal blood flow velocity was measured directly. In rabbits, volumetric renal blood flow was converted to velocity assuming a constant renal arterial diameter of 2.1 mm, consistent with prior reports (Chu *et al*., 2011). Blood density was assumed to be 1,050 kg/m^3^, and Savitzky-Golay filtering was performed on the acquired data series prior to analysis. Temporal variability in wave speed was quantified as the within-recording standard deviation of the beat-to-beat wave speed time series.

### Input Impedance Analysis

Input impedance was calculated using Fourier decomposition of the bilateral renal blood flow and arterial pressure waveforms (Figure S1). For the primary time-invariant analysis, individual cardiac cycles were linearly detrended, temporally normalized, ensemble-averaged, and then decomposed into the cardiac frequency and its harmonics. Coherence between pressure and flow showed that the first 8 harmonics maintained a coherence > 0.5 in rabbits and the first 9 maintained a coherence > 0.5 in swine, and thus 8 harmonics were calculated in rabbits and 9 harmonics in swine (Figure S2). At each harmonic, impedance modulus was calculated as the ratio of pressure amplitude to flow amplitude, and impedance phase was calculated as the phase difference between pressure and flow.

To characterize temporal variability, the same Fourier analysis was repeated on individual cardiac cycles to generate beat-to-beat time series of modulus and phase at each harmonic. Temporal variability was quantified as the within-recording standard deviation of each beat-to-beat impedance time series.

### Statistical Analysis

Data are presented as mean ± SD. Normality of model residuals was assessed using the Shapiro-Wilk test. Wave speed metrics were compared between innervated and surgically denervated kidneys using paired t-tests. Input impedance metrics were analyzed using repeated-measures analysis of variance with innervation status and harmonic as within-subject factors with Greenhouse-Geisser correction for sphericity. When the omnibus ANOVA identified a significant main effect of innervation or an innervation-by-harmonic interaction, post hoc comparisons between innervation states at individual harmonics were performed using the Holm-Sidak method to correct for multiple comparisons. For the stepwise catheter-denervation analysis, repeated-measures models incorporated denervation modality and ablation round as within-subject factors, with post hoc testing performed when justified by the omnibus model. Statistical significance was defined as a two-sided P < 0.05. Waveform analyses were performed in MATLAB R2025a (Mathworks, Natick, MA, USA), and statistical testing was performed in GraphPad Prism 11 (GraphPad Software, Boston, MA, USA).

## Results

### Wave Speed

Surgical renal denervation did not consistently affect wave speed (Figure 2). In conscious rabbits, a non-significant trend for a reduction in denervated kidneys (INV 19.0 ± 4.9, DNx 14.4 ± 5.5 m/s, P = 0.054) was observed, but this was not reproduced in swine (INV 19.2 ± 5.5, DNx 18.2 ± 4.3 m/s, P = 0.57). Beat-to-beat wave speed variability was unchanged by surgical denervation in both models. In swine undergoing unilateral catheter-based renal denervation, wave speed decreased similarly in both kidneys, indicative of a non-specific time-dependent influence on wave speed (P_ablation_ = 0.0064, P_denervation x ablation_ = 0.13, Figure 3). Consistent with this, multiplicity-corrected post-hoc testing showed bilateral renal arterial wave speed to be significantly decreased after 2, 3, and 4 rounds of ablation relative to baseline (P_adj_ = 0.049, P_adj_ = 0.0053, P_adj_ = 0.0064, respectively).

**Figure 1.**
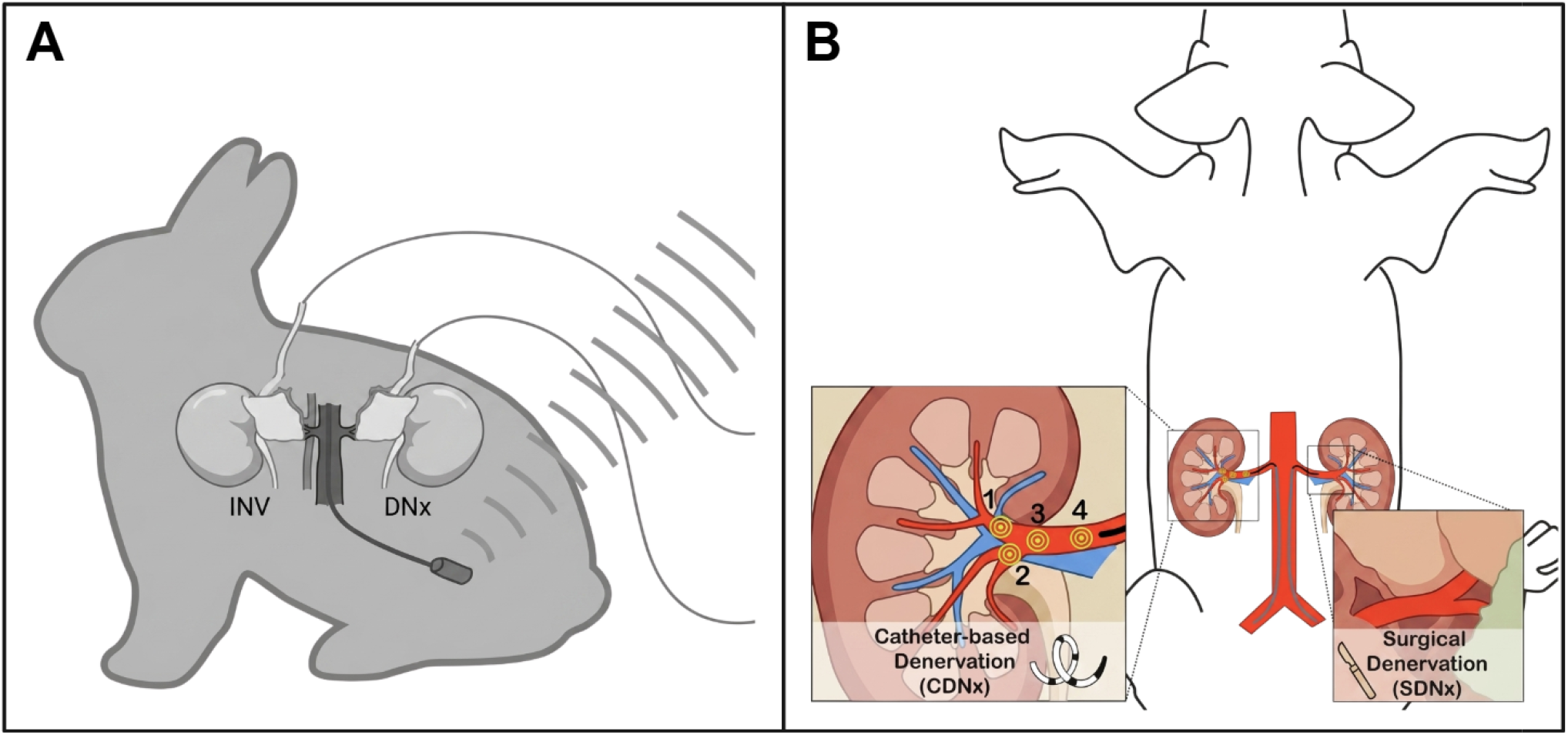
Preclinical renal denervation models. (A) Rabbits underwent unilateral surgical denervation and implantation of bilateral perivascular renal blood flow probes and an abdominal aortic pressure telemeter for subsequent conscious renal hemodynamic assessment and comparison to the innervated (INV) kidney. (B) Swine underwent unilateral surgical renal denervation (SDNx) one week prior to renal hemodynamic assessment via intravascular flow wires placed in the bilateral renal arteries using fluoroscopic guidance. After baseline measurements, swine underwent four rounds of catheter-based renal denervation (CDNx) in a distal-to-proximal fashion with hemodynamic assessment repeated between each round of ablation.

**Figure 2.**
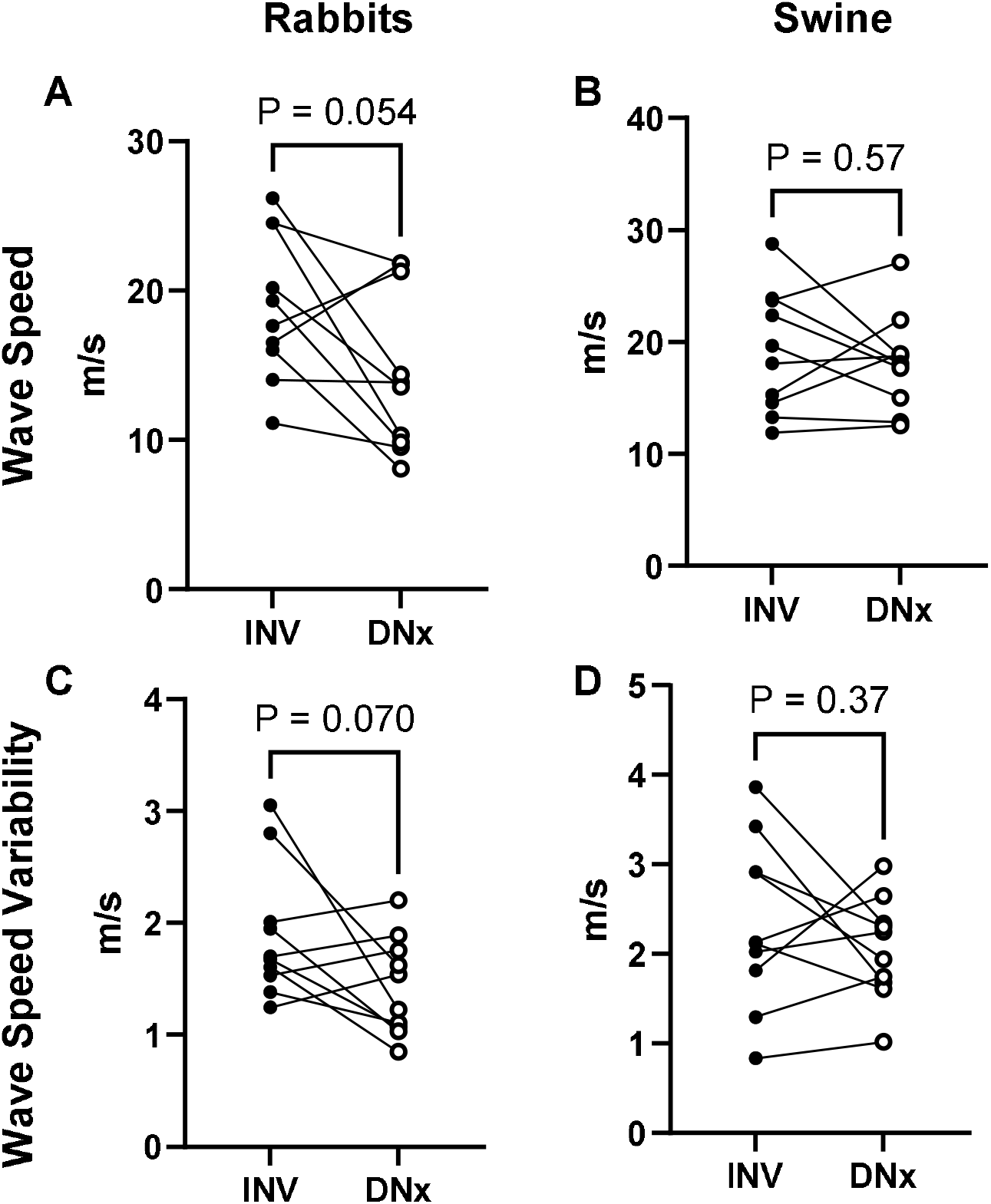
Renal arterial wave speed and wave speed variability in conscious rabbits and anesthetized swine after unilateral surgical renal denervation. Wave speed trends down in conscious rabbits (A) but not anesthetized swine (B). Wave speed variability also trends down in conscious rabbits (C) but not anesthetized pigs (D). All P values calculated from uncorrected paired t-tests, n = 10 per group for all analyses.

**Figure 3.**
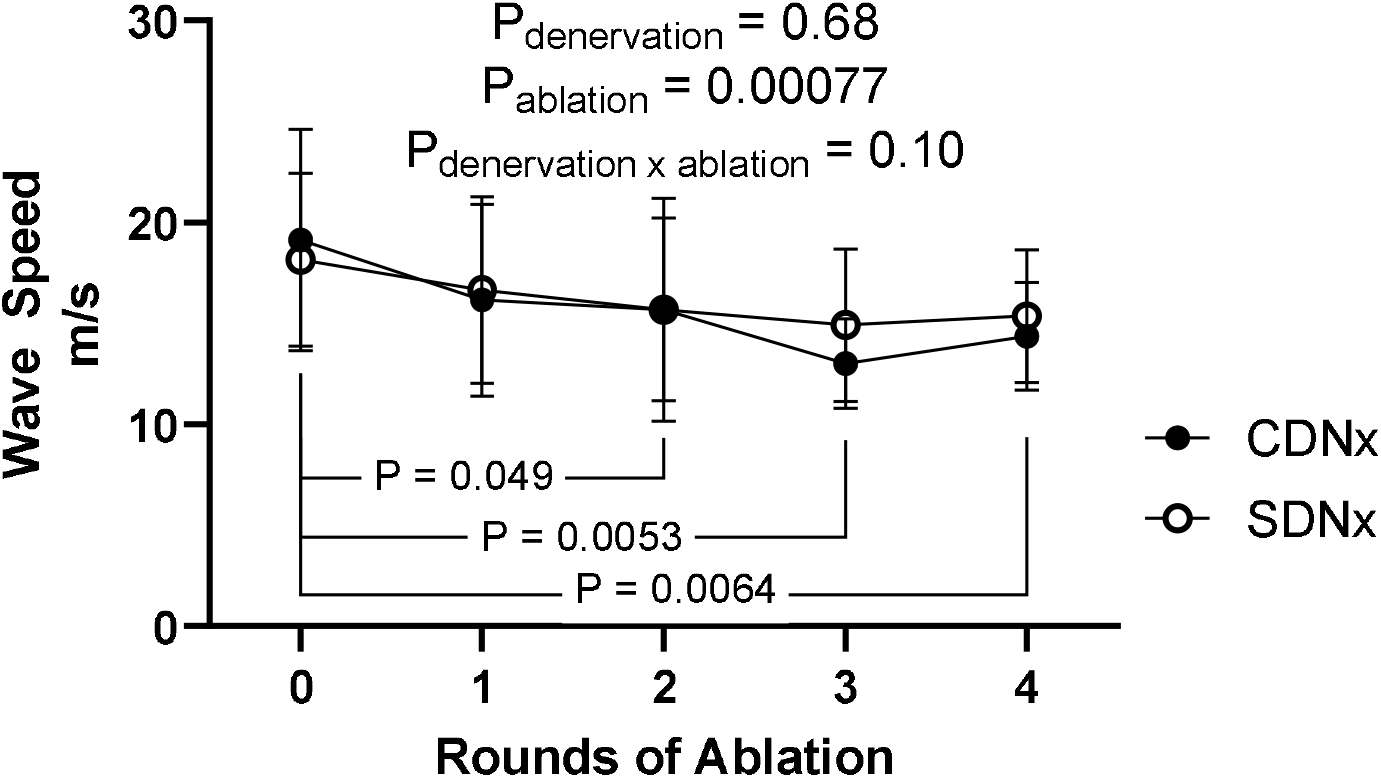
The effect of unilateral catheter-based renal denervation on wave speed. Wave speed decreases significantly in both renal arteries (P_ablation_ = 0.00077) with no augmentation by unilateral renal denervation (P_denervation x ablation_ = 0.10). P values calculated by repeated-measures ANOVA with denervation status and ablation round as within-subjects factors and post-hoc paired t-tests adjusted via the Holm-Sidak method for 10 comparisons, n = 10 per group for all analyses.

### Input Impedance

Contrary to our hypothesis, unilateral surgical renal denervation did not significantly affect renal arterial input impedance modulus in conscious rabbits (Figure 4A, 4B). Renal arterial phase shift was increased after unilateral denervation in rabbits in a harmonic-dependent manner (P_denervation_ = 0.0048, P_denervation x harmonic_ = 0.0060, Figure 4C). Multiplicity-corrected post-hoc testing found significant increases in phase shift of denervated kidneys at the 6^th^, 7^th^, and 8^th^ cardiac harmonics (P_adj_ = 0.00016, P_adj_ = 2.8e-6, P_adj_ = 3.1e-7, respectively). Conversely, unilateral denervation did not significantly alter phase shift between the innervated and surgically denervated kidney in our anesthetized swine model (Figure 4D).

**Figure 4.**
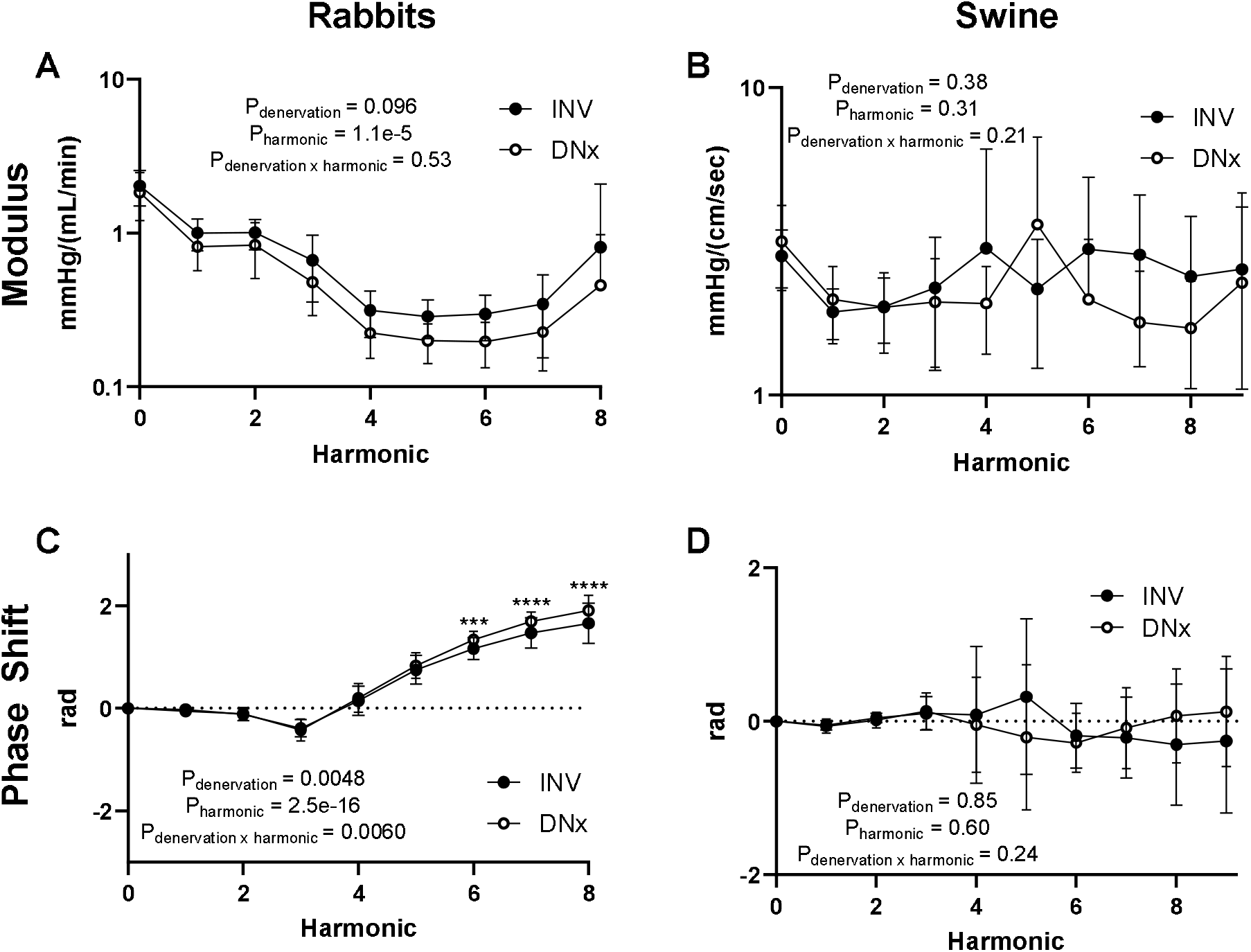
Renal arterial input impedance in conscious rabbits and anesthetized swine after unilateral surgical renal denervation. Surgical renal denervation did not significantly affect input impedance modulus in rabbits (A) or swine (B). Surgical renal denervation significantly increased phase shift in a harmonic-dependent fashion in rabbits (C) but not in swine (D). P values calculated by repeated-measures ANOVA with denervation status and harmonic as within-subjects factors with, in panel C, post-hoc paired t-tests adjusted via the Holm-Sidak method for 8 comparisons, ***, P_adj_ < 0.001, ****, P_adj_ < 0.0001, n = 10 per group for all analyses.

### Temporal Variability of Input Impedance

Unilateral surgical renal denervation did not produce a statistically significant effect on beat-to-beat variability of renal arterial input impedance modulus in rabbits or pigs (Figure 5A, 5B). While beat-to-beat variability of phase shift was not significantly affected in rabbits (Figure 5C), unilateral surgical denervation decreased phase shift variability in swine (P_denervation_ = 0.0048, Figure 5D). Post-hoc testing in swine identified higher phase shift variability in innervated kidneys at the 7^th^ and 8^th^ harmonics compared to surgically denervated kidneys (P_adj_ = 0.036, P_adj_ = 0.036, respectively).

**Figure 5.**
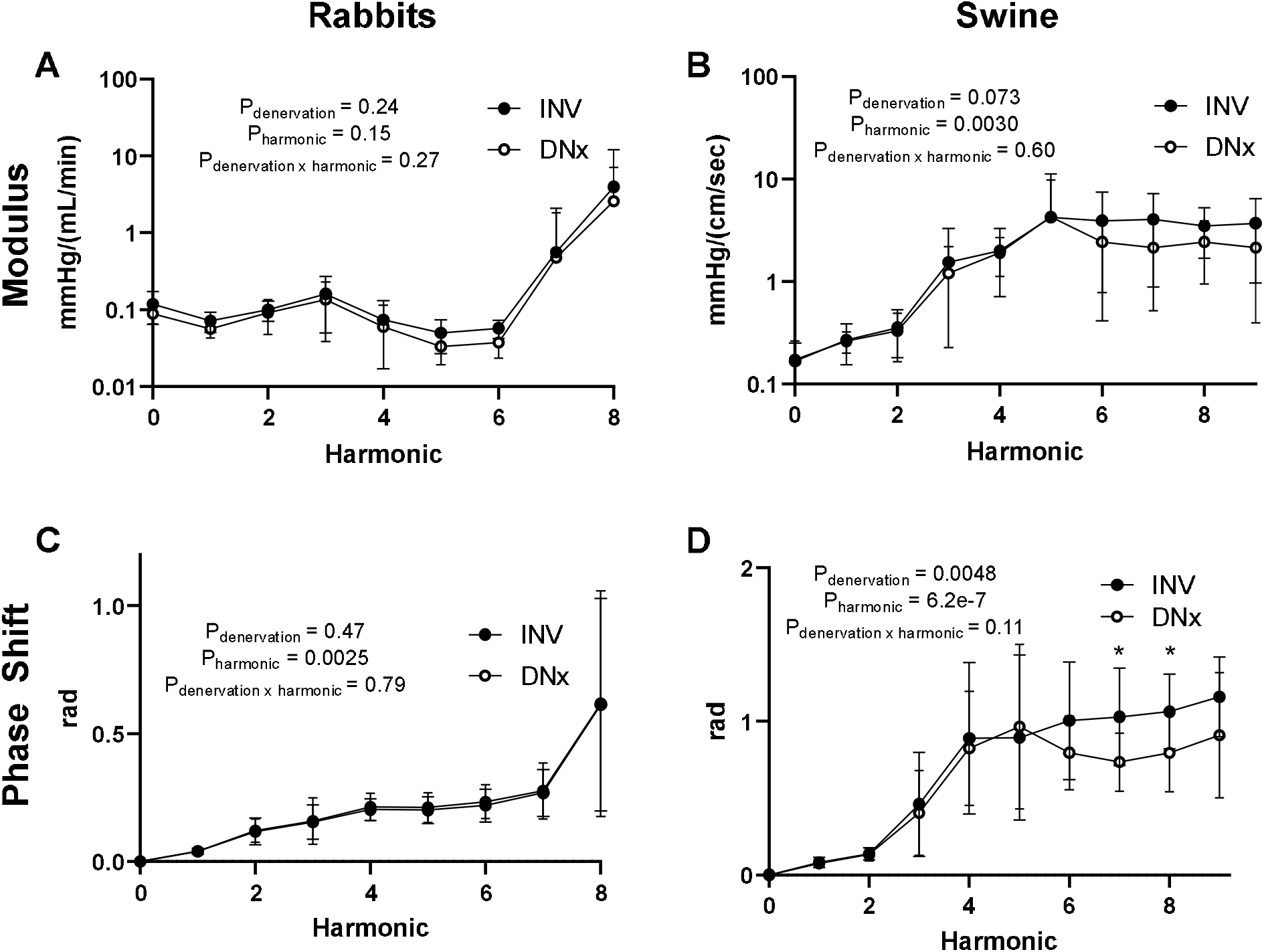
Renal arterial input impedance variability in conscious rabbits and anesthetized swine after unilateral surgical renal denervation. Surgical renal denervation did not significantly affect beat-to-beat renal arterial impedance modulus variability in rabbits (A) or swine (B). Although no significant effect was observed in rabbits (C), surgical renal denervation decreased renal arterial impedance phase shift variability in pigs (D). P values calculated by repeated-measures ANOVA with denervation status and harmonic as within-subjects factors with, in panel D, post-hoc paired t-tests for INV vs. DNx adjusted via the Holm-Sidak method for 9 comparisons, *, P_adj_ < 0.05, n = 10 per group for all analyses.

However, this between-kidney difference was eliminated after one round of catheter-based renal nerve ablation involving only a single renal branch artery (P_denervation_ = 0.42, Figure 6A) with no effect of subsequent rounds of ablation (Figure 6B-D).

**Figure 6.**
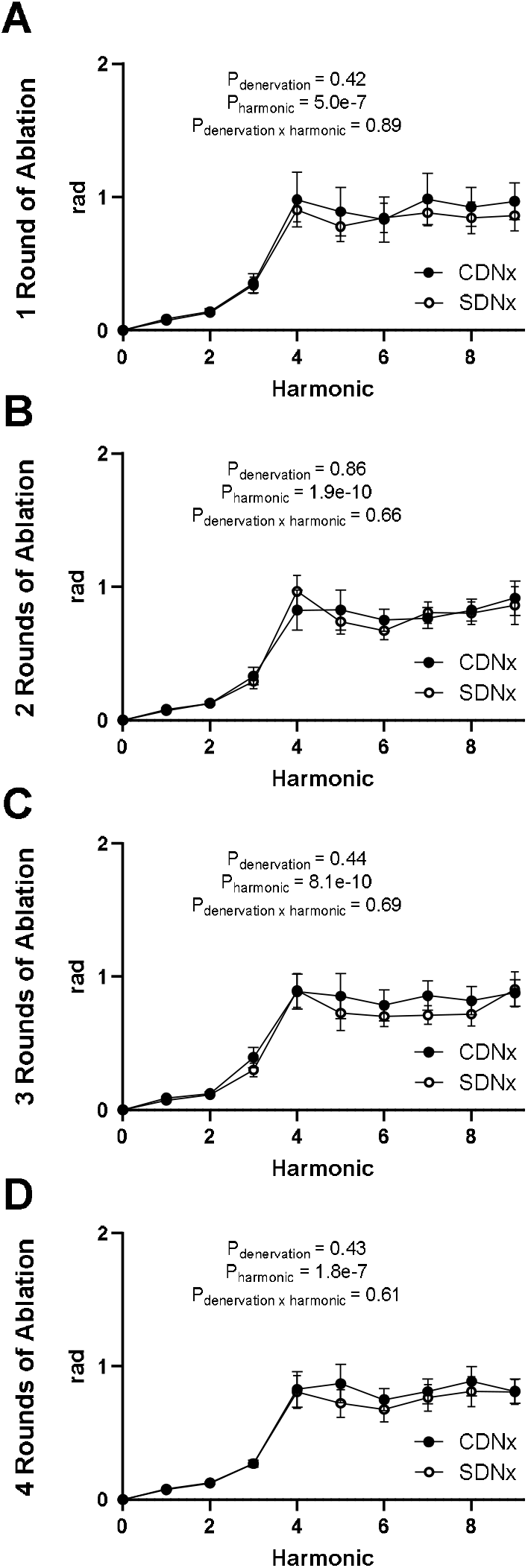
The effect of unilateral catheter-based renal denervation on phase shift variability in swine. After one round of catheter-based ablation, the baseline between-kidney difference in renal arterial input impedance phase shift variability is abolished (A), and additional rounds of catheter-based denervation have no additional effect on this measure (B-D). P values calculated by repeated-measures ANOVA with denervation status and ablation round as within-subjects factors, n = 10 per group for all analyses.

## Discussion

A major barrier to renal denervation is the lack of biomarkers to guide patient selection beforehand and confirm effective denervation during the procedure. In this study, we tested two potential biomarkers of sympathetic innervation, wave speed and input impedance, and their beat-to-beat variability in conscious rabbits and anesthetized swine. In these two complementary preclinical models, no metric showed the species-spanning, dose-dependent behavior expected of a direct biomarker of renal sympathetic innervation. Wave speed was unchanged by surgical denervation in swine, showed only a borderline reduction in rabbits, and decreased similarly in the previously surgically denervated kidney while the contralateral kidney underwent catheter-based denervation. Renal input impedance phase shift was increased in rabbits in a harmonic-dependent manner in rabbits but not in swine. Input impedance phase shift time variability was decreased after surgical renal denervation in swine but not in rabbits. Moreover, this significant difference was eliminated after radiofrequency ablation of the largest branch artery with the remaining branch artery (or arteries) still innervated, indicating an inability to reflect progressive denervation. Taken together, these findings argue against using these measures in their present form as direct physiological surrogates of renal sympathetic innervation or outflow.

The models used in this manuscript provide powerful insight into renal sympathetic control. In both models, simultaneous recordings of pressure and blood flow were obtained from kidneys differing only in innervation that were exposed to the same perfusion pressure, neurohumoral milieu, and, in swine, anesthetic depth. The use of surgical renal denervation, the gold-standard research denervation modality, enhances confidence in the innervation difference between kidneys, and the use of a second-generation clinically approved catheter-based system employed in a stepwise fashion allowed for screening of dose-responsiveness between these candidate markers for clinically relevant use. We have previously validated renal denervation in these same cohorts using functional and molecular testing, and alternative forms of pressure-flow analysis have revealed significant, dose-dependent, cross-species differences (Pellegrino *et al*., 2020, 2025).

These models also differ in ways that impact interpretation of the results. Rabbits and swine have very different cardiac frequencies (3.60 ± 0.28 Hz in rabbits compared to 1.19 ± 0.19 Hz in swine), and cardiac frequency is the foundation of input impedance analysis. Thus, while the harmonics analyzed for input impedance computations were similar, the absolute frequencies they represent differed approximately threefold, making direct cross-species comparisons challenging. In addition, rabbits were studied in the conscious resting state after many sessions of acclimation to a procedure room while swine were anesthetized with a ketamine and midazolam infusion, which certainly affects autonomic outflow, albeit less than volatile anesthetics. The instrumentation used to measure blood flow also differed substantially. In rabbits, perivascular transit-time flow probes yielded volumetric renal blood flow measurements; in swine, intravascular Doppler wires measured renal blood flow velocity. While perivascular transit-time flow probes produce high-fidelity flow measurements, Doppler flow velocity represents the clinically viable method for measuring renal blood flow. These differences provide strength to assess the generalizability of arterial hemodynamic measures as a surrogate for renal sympathetic innervation across species, heart rate, anesthesia, and instrumentation.

Prior clinical reports have described associations between arterial mechanics and blood pressure response after renal denervation (Finegold *et al*., 2016). Those observations were made in patients with treatment-resistant hypertension who likely exhibit extensive vascular remodeling and chronic sympathetic activation. In that setting, arterial hemodynamic properties may be useful predictors of clinical response without necessarily serving as direct physiological biomarkers of renal sympathetic innervation. Our results therefore should not be interpreted as disproving the previously reported predictive association. Rather, they suggest that the biological information carried by wave speed in hypertensive patients may extend beyond the degree of renal sympathetic innervation itself.

A range of approaches has been explored to provide physiological feedback for renal denervation. Our group previously showed that renal sympathetic vasomotion, a time-varying pressure-flow measure of sympathetic vascular control, decreases with both surgical denervation and catheter-based renal denervation and demonstrates a dose-response relationship during stepwise radiofrequency ablation (Pellegrino *et al*., 2020, 2025). Other demonstrated approaches include renal norepinephrine spillover and evoked responses to renal nerve stimulation (Esler *et al*., 2010; De Jong *et al*., 2018).

While renal norepinephrine spillover remains the gold-standard method for clinical assessment of regional renal sympathetic activity, its use requires specialized invasive sampling and offline tracer-based analysis, making it a research technique unsuitable for widespread clinical deployment. Renal sympathetic nerve stimulation maps electrically evoked physiological responses to guide denervation therapy but cannot facilitate patient selection or clearly confirm denervation completeness. The practical limitations of these other approaches motivate interest in passive pressure-flow biomarkers that can be derived either non-invasively via transabdominal ultrasound or as part of catheter-based minimally invasive procedures to guide renal denervation.

Our study had several limitations. First, the experiments were performed in healthy animals rather than in models of chronic hypertension. This limits direct generalization to the diseased vasculature of patients undergoing renal denervation, particularly because hypertension itself alters arterial stiffness, vascular remodeling, and sympathetic tone. At the same time, the absence of induced cardiovascular disease allowed us to test the relationship between sympathetic innervation and arterial mechanics with fewer disease-related confounders and less between-animal heterogeneity. Second, time-dependent effects were observed in anesthetized swine independent of catheter-based renal denervation as shown in Figure 3. Despite efforts to control these aspects of the experiment, this could reflect changes in anesthetic depth, volume status, and neurohumoral activation. Third, we evaluated a limited set of arterial hemodynamic measures based on prior studies. Other arterial indices or analyses of the temporal structure of these measurements may prove more sensitive to sympathetic vascular control.

In conclusion, renal arterial wave speed, input impedance, and their beat-to-beat variability did not consistently reflect renal sympathetic innervation in tightly controlled rabbit and swine models and did not demonstrate the progressive response expected during stepwise catheter-based denervation. These findings make them poor candidates, in their present form, for direct assessment of renal innervation and thereby renal sympathetic outflow and denervation completeness. This does not exclude the possibility that arterial hemodynamic properties may predict clinical response in patients with hypertension, nor does it exclude the possibility that more advanced dynamic analyses of renal arterial mechanics might better reflect sympathetic vascular control. Defining a practical physiological biomarker of sympathetic outflow remains an important challenge for the field of renal denervation.

## Data Availability Statement

Data and code are available on figshare (doi: 10.6084/m9.figshare.33335151). The corresponding author is responsible for maintenance of this data repository.

## Ethics Statement

All animal experiments were reviewed and approved by the Institutional Animal Care and Use Committee of the University of Nebraska Medical Center and were conducted in accordance with the NIH Guide for the Care and Use of Laboratory Animals.

## Author Contributions

CD contributed to conceptualization, formal analysis, methodology, visualization, and writing of the original draft. AMS contributed to conceptualization, funding acquisition, investigation, methodology, project administration, resources, supervision, and reviewing and editing of the manuscript. IHZ contributed to conceptualization, funding acquisition, investigation, project administration, resources, supervision, and reviewing and editing of the manuscript. YSC contributed to conceptualization, funding acquisition, investigation, methodology, project administration, resources, and reviewing and editing of the manuscript. HJW contributed to conceptualization, funding acquisition, investigation, methodology, project administration, resources, and reviewing and editing of the manuscript. PRP contributed to conceptualization, data curation, formal analysis, funding acquisition, investigation, methodology, project administration, resources, software, supervision, validation, visualization, and writing of the original draft.

## Figure Legends

**Supplemental Figure 1.**
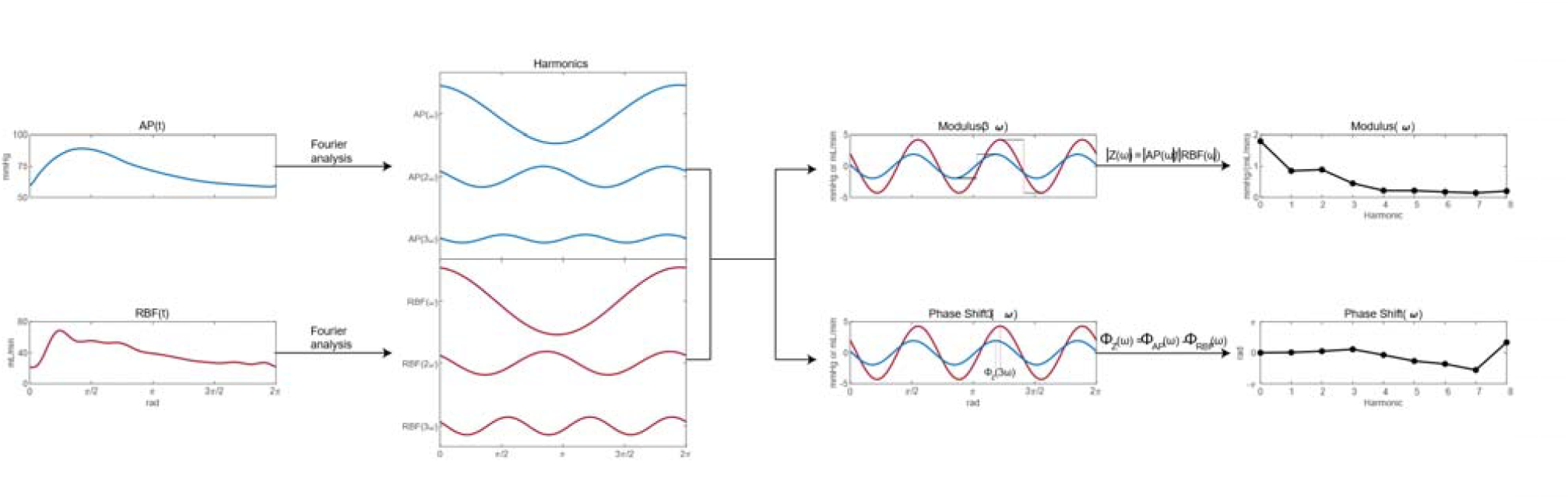
Ensemble-averaged and beat-to-beat input impedance analysis workflow.

**Supplemental Figure 2.**
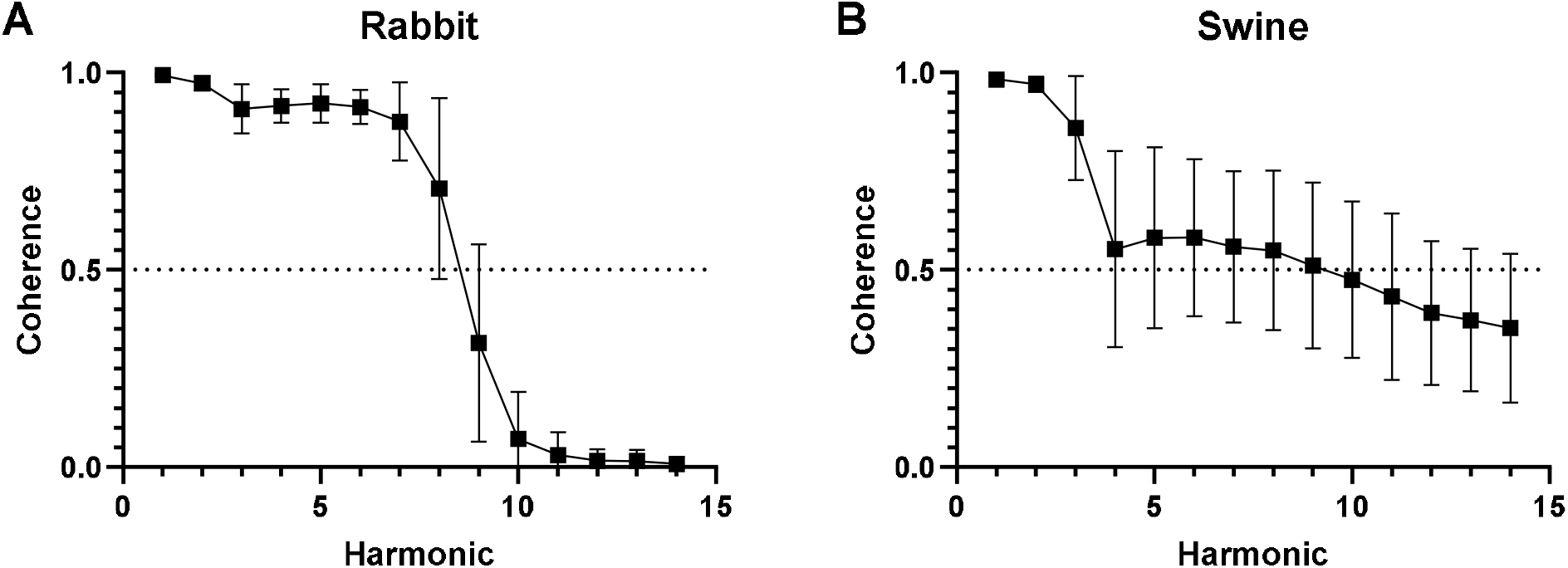
Coherence as a function of harmonic. Pressure-flow coherence remains above 0.5 for the first 8 harmonics in rabbits (A) and the first 9 harmonics in pigs (B). Data derived from all 20 simultaneous pressure-flow recordings in rabbits and all 100 simultaneous pressure-flow recordings in swine.

## References

Azizi, M. et al. (2018) “Endovascular ultrasound renal denervation to treat hypertension (RADIANCE-HTN SOLO): a multicentre, international, single-blind, randomised, sham-controlled trial,” The Lancet, 391(10137), pp. 2335–2345. Available at: 10.1016/S0140-6736(18)31082-1.

Azizi, M. et al. (2022) “Effects of Renal Denervation vs Sham in Resistant Hypertension After Medication Escalation: Prespecified Analysis at 6 Months of the RADIANCE-HTN TRIO Randomized Clinical Trial,” JAMA Cardiology, 7(12), pp. 1244–1252. Available at: 10.1001/JAMACARDIO.2022.3904.

Böhm, M. et al. (2020) “Efficacy of catheter-based renal denervation in the absence of antihypertensive medications (SPYRAL HTN-OFF MED Pivotal): a multicentre, randomised, shamcontrolled trial.,” Lancet (London, England), 395(10234), pp. 1444–1451. Available at: 10.1016/S0140-6736(20)30554-7.

Carlström, M., Wilcox, C.S. and Arendshorst, W.J. (2015) “Renal Autoregulation in Health and Disease.,” Physiological reviews, 95(2), pp. 405–511. Available at: 10.1152/physrev.00042.2012.

Chu, Y. et al. (2011) “The morphology and haemodynamics of the rabbit renal artery: evaluation by conventional and contrast-enhanced ultrasonography,” Laboratory animals, 45(3), pp. 204–208. Available at: 10.1258/LA.2011.011022.

Davies, J.E. et al. (2006) “Use of simultaneous pressure and velocity measurements to estimate arterial wave speed at a single site in humans,” American journal of physiology. Heart and circulatory physiology, 290(2). Available at: 10.1152/AJPHEART.00751.2005.

Davies, J.E. et al. (2017) “Use of the Instantaneous Wave-free Ratio or Fractional Flow Reserve in PCI,” The New England journal of medicine, 376(19), pp. 1824–1834. Available at: 10.1056/NEJMOA1700445.

DiBona, G.F. and Kopp, U.C. (1997) “Neural control of renal function,” Physiological Reviews, 77(1), pp. 75–197.

Esler, M. (2014) “Illusions of truths in the Symplicity HTN-3 trial: generic design strengths but neuroscience failings.,” Journal of the American Society of HypertensionJ: JASH, 8(8), pp. 593–8. Available at: 10.1016/j.jash.2014.06.001.

Finegold, J. et al. (2016) “Systematic evaluation of haemodynamic parameters to predict haemodynamic responders to renal artery denervation,” Presented at EuroPCR 2016. [Abstract].

Foss, J.D., Fink, G.D. and Osborn, J.W. (2016) “Differential role of afferent and efferent renal nerves in the maintenance of early- and late-phase Dahl S hypertension,” American journal of physiology. Regulatory, integrative and comparative physiology, 310(3), pp. R262–R267. Available at: 10.1152/AJPREGU.00408.2015.

Guild, S.J. et al. (2001) “Dynamic relationship between sympathetic nerve activity and renal blood flow: a frequency domain approach.,” American journal of physiology. Regulatory, integrative and comparative physiology, 281(1), pp. R206–12. Available at: http://www.ncbi.nlm.nih.gov/pubmed/11404295 (Accessed: May 19, 2016).

Kiuchi, M.G. et al. (2019) “Renal Denervation Update From the International Sympathetic Nervous System Summit: JACC State-of-the-Art Review,” Journal of the American College of Cardiology, 73(23), pp. 3006–3017. Available at: 10.1016/J.JACC.2019.04.015.

Mahfoud, F. et al. (2022) “Long-term efficacy and safety of renal denervation in the presence of antihypertensive drugs (SPYRAL HTN-ON MED): a randomised, sham-controlled trial,” The Lancet, 399(10333), pp. 1401–1410. Available at: 10.1016/S0140-6736(22)00455-X.

Mitchell, G.F. et al. (2010) “Arterial stiffness and cardiovascular events: the Framingham Heart Study,” Circulation, 121(4), pp. 505–511. Available at: 10.1161/CIRCULATIONAHA.109.886655.

O’Rourke, M.F. (1982) “Vascular impedance in studies of arterial and cardiac function.,” 10.1152/physrev.1982.62.2.570, 62(2), pp. 570–623. Available at: 10.1152/PHYSREV.1982.62.2.570.

Pace, J.B. (1971) “Sympathetic control of pulmonary vascular impedance in anesthetized dogs,” Circulation research, 29(5), pp. 555–568. Available at: 10.1161/01.RES.29.5.555.

Paton, J.F.R. et al. (2026) “Multimodal, device-based therapeutic targeting of the cardiovascular autonomic nervous system,” Nature reviews. Cardiology, 23(4), pp. 255–278. Available at: 10.1038/S41569-025-01212-4.

Pellegrino, P.R. et al. (2020) “Quantification of Renal Sympathetic Vasomotion as a Novel End Point for Renal Denervation,” Hypertension (Dallas, Tex. J: 1979), 76(4), pp. 1247–1255. Available at: 10.1161/HYPERTENSIONAHA.120.15325.

Pellegrino, P.R. et al. (2025) “Sympathetic Vasomotion Reflects Catheter-Based Radiofrequency Renal Denervation,” Hypertension, 82(7), pp. 1261–1270. Available at: 10.1161/HYPERTENSIONAHA.125.24980,.

Saxena, M. et al. (2022) “Predictors of blood pressure response to ultrasound renal denervation in the RADIANCE-HTN SOLO study,” Journal of human hypertension, 36(7), pp. 629–639. Available at: 10.1038/S41371-021-00547-Y.

Schiller, A.M., Pellegrino, P.R. and Zucker, I.H. (2016) “Renal nerves dynamically regulate renal blood flow in conscious, healthy rabbits.,” American journal of physiology. Regulatory, integrative and comparative physiology, 310(2), pp. R156-66. Available at: 10.1152/ajpregu.00147.2015.

Scott-Solomon, E., Boehm, E. and Kuruvilla, R. (2021) “The sympathetic nervous system in development and disease,” Nature reviews. Neuroscience, 22(11), pp. 685–702. Available at: 10.1038/S41583-021-00523-Y.

